# SARS-COV-2 RBD Oral Vaccine Boosted by Mucosal Immune Adjuvant LTB26 via DCs and B Cells Activation in Mice

**DOI:** 10.1101/2020.04.06.025981

**Authors:** Yongping Ma, Qinlin Shi, Qiujuan Wang, Sijing Chen, Sijie Gan, Changyin Fang, Yanxi Shen, Min Jiang, Tao Lin, Fangzhou Song

## Abstract

Although several SARS-COV-2 vaccines have been approved, no one oral live vaccine is available. Here, an oral SARS-COV-2 RBD live vaccine containing LTB26 adjuvant has been developed. BALB/c mice are oral vaccinated with attenuated *Salmonella typhimurium* SL7207 containing pcDNA3.1-LTB26RBD or pcDNA3.1-RBD plasmids. The result shows that the high level of RBD specific antibody is produced in pcDNA3.1- LTB26RBD treatment. The mechanism indicates that LTB26 enhances RBD antibody production by significantly upregulating the activity of MHC II^+^ DCs and CD19^+^CD45^+^ B cells. LTB26 mutant is derived from heat-labile enterotoxin B subunit (LTB) wild type of *Escherichia coli* with enhanced immune adjuvanticity. Based on the pre-experiment result that SL7207 interferes the function of LTB26, the purified LTB26 was mixed with purified human rotavirus VP8 antigen to explore the mechanism of adjuvant. The results suggests that LTB26 enhances mucosal immune responses via increased of BCR and MHC II^+^ expression. Furthermore, LTB26 promotes both Th1 and Th2 cell mediated immunity. Therefore, LTB26 maybe a potent adjuvant for mucosal vaccine development in view of the safety of LTB26 than LT toxin.

## Introduction

Since December 2019, the 2019 novel coronavirus (2019-nCoV, SARS-COV-2) has infected over 95.38 million cases worldwide with about 2.03 million deaths. Although about 120 vaccines are in different stage of clinical trials and some inactivated or mRNA vaccine candidates have been approved, little is oral vaccination. Therefore, safe and effective mucosal vaccines are still necessary to prevent infection from SARS-COV-2.(Poland, Ovsyannikova et al., 2020) SARS-COV-2 cell entry depends on ACE2-RBD interaction(Hoffmann, Kleine-Weber et al., 2020). Therefore, receptor binding domain (RBD) of SARS-COV-2 spike protein (S protein) is the first target for vaccine development. As a pathogen of mucosal infection, mucosal immunity is very important to prevent of SARS-COV-2.

LTB26 is derived from heat-labile enterotoxin B subunit (LTB) of *Escherichia coli* with enhanced mucosal immune adjuvanticity. From the perspective of clinical application, LTB and its derivatives are safe without toxicity than LT toxin(da Hora, Conceicao et al., 2011). LT belongs to the members of the AB5 bacterial toxin family and consists of five B subunit (LTB) with a single catalytically active A subunit (LTA)(Merritt, Zhang et al., 2002). LTA has toxic effects of ADP-ribosylating activity and LTB binds to the cell membrane with GM1 ganglioside (GM1) following delivery LTA into cells inducing infectious diarrhea(da Hora et al., 2011). LT has been reported as mucosal immune adjuvant in 1988(Clemens, Sack et al., 1988). However, it fails to meet the clinical requirement for its toxicity(da Hora et al., 2011, Ma, 2016). Nevertheless, LTB is also not applicable for weaker adjuvanticity comparing with LT. Therefore, the construction of non-toxic LT has been a research hotspot in recent years(Clements & Norton, 2018, Ghazali-Bina, Pourmand et al., 2019, Ma, 2016, Nashar, 2014, White, Haghighi et al., 2017). However, there is no report to develop adjuvanticity-enhanced LTB mutant.

In this study, we optimized an adjuvanticity-enhanced LTB26 mutant from four LTB mutants (Holmner, Mackenzie et al., 2011, Ma, 2016). By fusion expressing with LTB26 adjuvant, a promising live SARS-COV-2 LTB26RBD mucosal vaccine was constructed using attenuated *Salmonella typhimurium* SL7207 to boost systematic immune responses.

## Results

### LTB26-RBD recombinant was constructed successfully

After commercial synthesis of LTB26-RBD and RBD sequences, pcDNA3.1-LTB26RBD and pcDNA3.1-RBD plasmids were constructed, respectively. Then attenuated *Salmonella typhimurium* SL7207 (SL7207) was electrically transformed with pcDNA3.1-LTB26RBD, pcDNA3.1-RBD and pcDNA3.1 plasmids at 2.5kv and the restriction enzymes analysis confirmed the recombinants of LTB26-RBD and RBD vaccine candidates (Fig. 1A).

**Figure 1.**
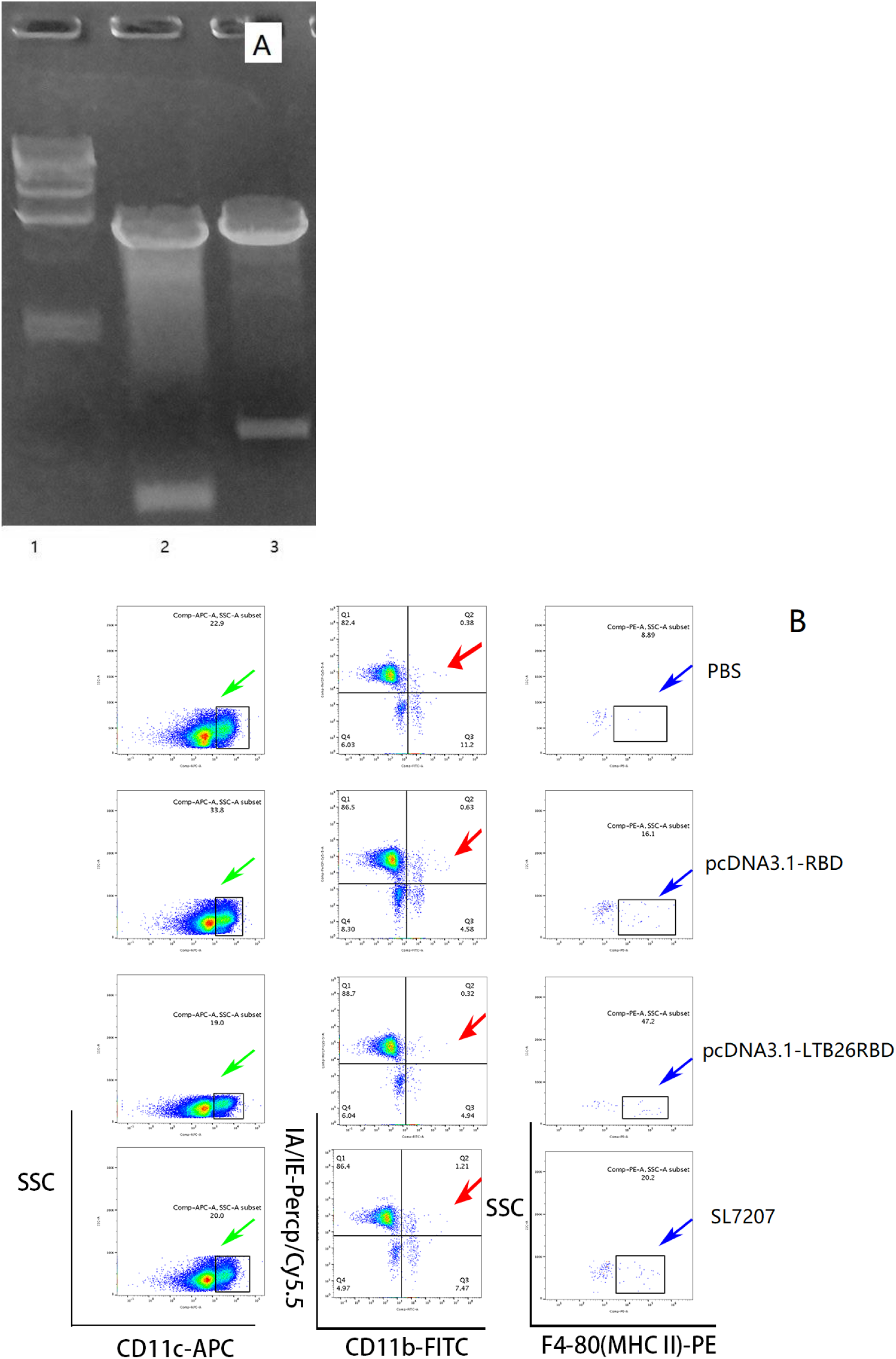
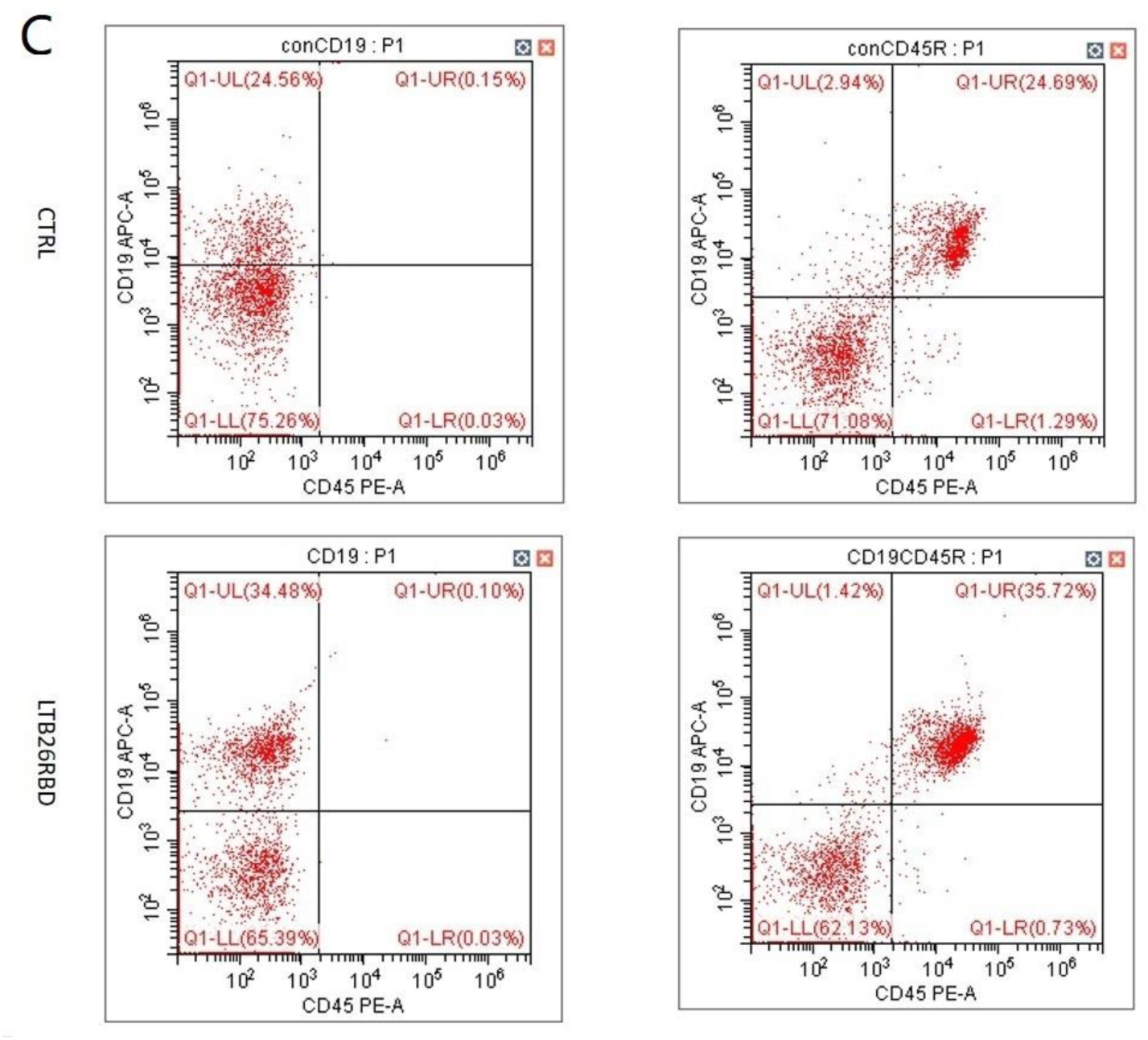
FCM analysis of dendritic cells (DCs) and B cells. (A) pcDNA3.1-LTB26-RBD analyzed with restriction enzymes. Lane 1: DNA marker, lane 2: pcDNA3.1- RBD, lane 2: pcDNA3.1-LTB26-RBD. (B) LTB26 significantly activated MHC II^+^ DCs and increased about 29.3% proportion than that of RBD alone vaccination. There were no significant differences between Salmonella and Salmonella-RBD vaccination mice. (C) LTB26 significantly activated CD19^+^CD45^+^ B cells and increased about 14.5% proportion than that of RBD alone vaccination.

### LTB26 boosted immune responses via upregulating MHC II+ DCs

In order to decipher the mechanism, dendritic cells (DCs) and B cells from vaccinated mice were purified and analyzed with FCM (flow cytometry). The results suggested that LTB26 activated MCH II^+^ DCs than *Salmonella typhimurium* SL7207 (SL7207) or RBD alone. The ratio of LTB26RBD to RBD was 29.3 times, LTB26RBD to PBS was 53.1 times, and LTB26RBD to SL7207 was 23.4 times. However, The ratio of SL7207 to PBS was 22.7 times. Even though there were no significant MCH II^+^ DCs differences between SL7207 and SL7207-RBD vaccination mice comparing with SL7207-LTB26RBD treatment. However, SL7207-RBD and SL7207 vaccinations were significantly upregulated DCs than that of PBS treatment (Fig. 1B). That meant, LTB26 adjuvant was necessary for *Salmonella*-based vaccine construction. Meanwhile, LTB26 significantly activated CD19^+^CD45^+^ B cells than PBS treatment and increased about 14.5% proportion (Fig. 1C). As a summary, LTB26 significantly upregulated both DCs and B cells activity to enhance antigen presentation and antibody production.

### LTB26 enhanced mice to produce high-level IgG antibody

After the 3^rd^ boosting vaccination, BALB/c mice produced high-level RBD specific antibody in SL7207-LTB26RBD immune group (Fig. 2, p<0.025). The result indicated that LTB26 has promising adjuvant activity. However, we did not continue the vaccination of SL7207-RBD single group for the absent of LTB26 adjuvant according to the primary test (Fig. 1B, C).

**Figure 2.**
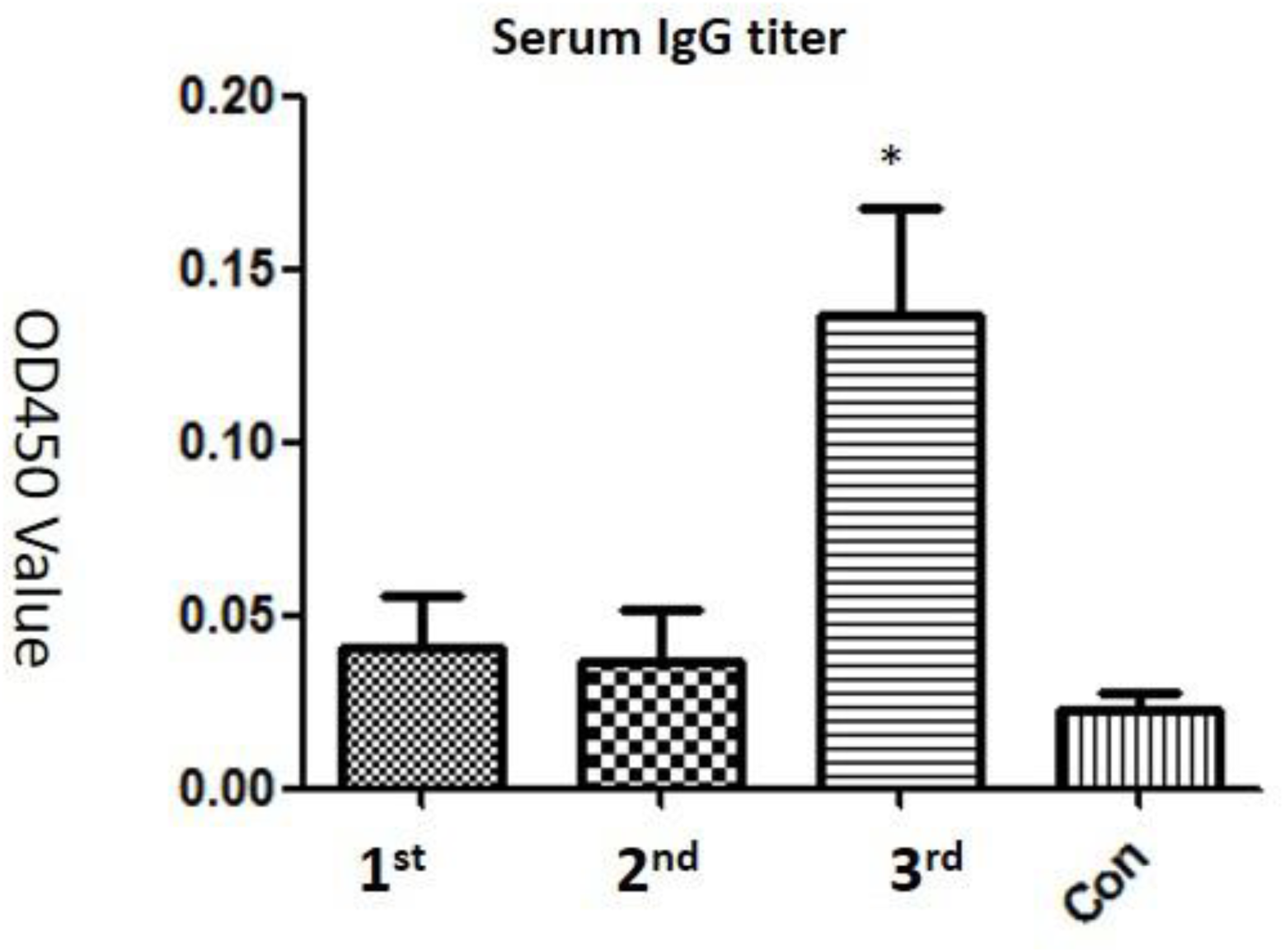
Immune responses analysis. Serum IgG titer on post-vaccination of 10 d, 20 d and 30 d, respectively; *, p<0.02. PBS treatment was used as control group.

### The characteristic of LTB mutants

The pET32a-LTB, pET32a-LTB26, pET32a-LTB34, pET32a-LTB57 and pET32a-LTB85 were confirmed by sequencing and the products of the five proteins were detected by SDS-PAGE. Then, the proteins were purified using BeaverBeads™ His-tag protein purification kit (Beaverbio, Suzhou, China) and were detected a band of 30 KD protein using SDS-PAGE (Fig.3A). The protein concentration of LTB was 3.412 µg/µl, LTB26, 3.215 µg/µl, LTB34, 3.325 µg/µl, LTB57, 3.011 µg/µl, and LTB85, 1.375 µg/µl, respectively.

**Figure 3.**
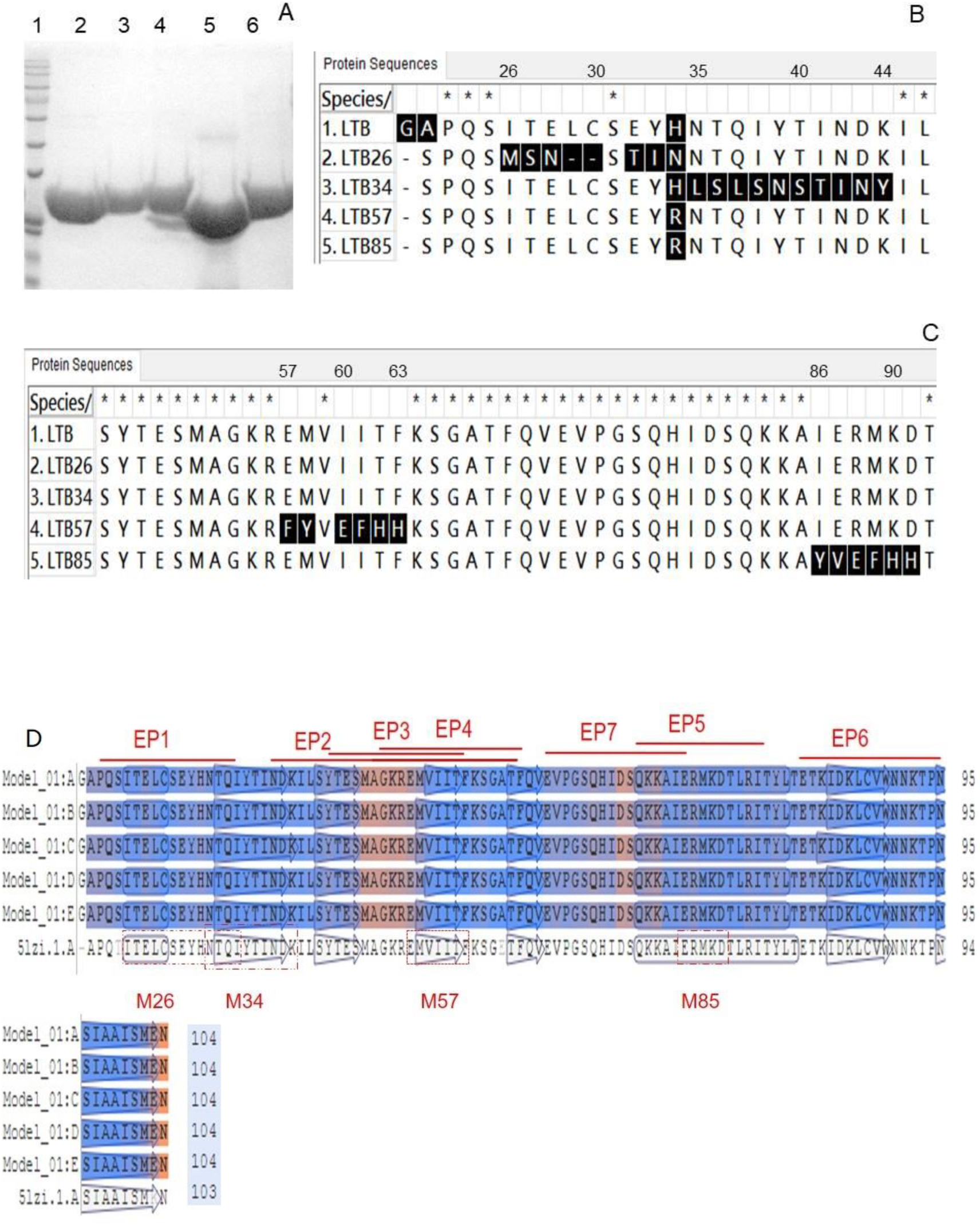
The construction and analysis of LTB mutants. (A) LTB and the mutants were purified and detected in SDS-PAGE. Lane 1: standard proteins; lane 2-6: LTB, LTB26, LTB34, LTB57, and LTB85; (B) Alignment of LTB26 and LTB34 with LTB; (C) alignment of LTB57 and LTB85 with LTB. The mutations were highlighted with black background; (D) B cell and T cell epitopes of LTB and the LTB mutants regions located in LTB epitopes.

Compared with LTB, LTB26 was mutated from I26T27E28L29C30 to M26S27N28 and from E32Y33H34 to T32I33N34, and meanwhile deleted L29C30 residues (Fig.3B). In turn, LTB34 was mutated from N35T36 Q37I38Y39T40I41N42D43K44 to L35S36L37S38N39S40T41I42N43 Y44 (Fig.1B), LTB57 was mutated from E57M58 to F57Y58 and from I60I61T62F63 to E60F61H62H63 (Fig.3C) and LTB85 was mutated from I86E87R88M89K90D91 to Y86V87E88F89H90H91, respectively (Fig.3C). As a summary, the mutations were located in epitope 1(EP1) of LTB (LTB26), middle EP1 and EP2 (LTB34), EP2,3,4 (LTB57) and EP5 (LTB85), respectively (Fig.3D)(Takahashi, Kiyono et al., 1996).

### LTB26 significantly enhanced both systematic and mucosal immune response

Because *Salmonella* SL7207 interfered with the adjuvant activity of LTB26 as previously described (Figure 2A), we used the purified human rotavirus VP8 antigen (hRV VP8 or VP8) to study the molecular mechanism of the adjuvant of purified LTB26 and other mutants. The serum VP8 specific IgG levels of LTB+VP8, LTB26+VP8 and LTB34+VP8 were significantly higher than that of the VP8 alone group (Fig.4A, p<0.01). However, the serum IgG level of LTB34+VP8 vaccination was slightly higher than that of LTB+VP8 treatment, but lower than that of LTB26+VP8 vaccination (Fig.4A, p<0.01). On the contrary, the serum IgG levels of LTB57+VP8 and LTB85+VP8 vaccinations were lower than that of LTB+VP8 groups (Fig. 4A). Similarly, the variation trends of lung mucosal VP8 specific sIgA were consistent with that of serum IgG among the LTB+VP8 and the other LTB mutants vaccination on 21 d post-vaccination (p<0.001) (Fig. 4B). LTB26 also elicited the highest mucosal immune response compared with LTB and LTB34, respectively. Thus, LTB26 was the optimal adjuvant mutant of LTB to elicit robust systematic and mucosal immune responses in this study.

**Figure 4.**
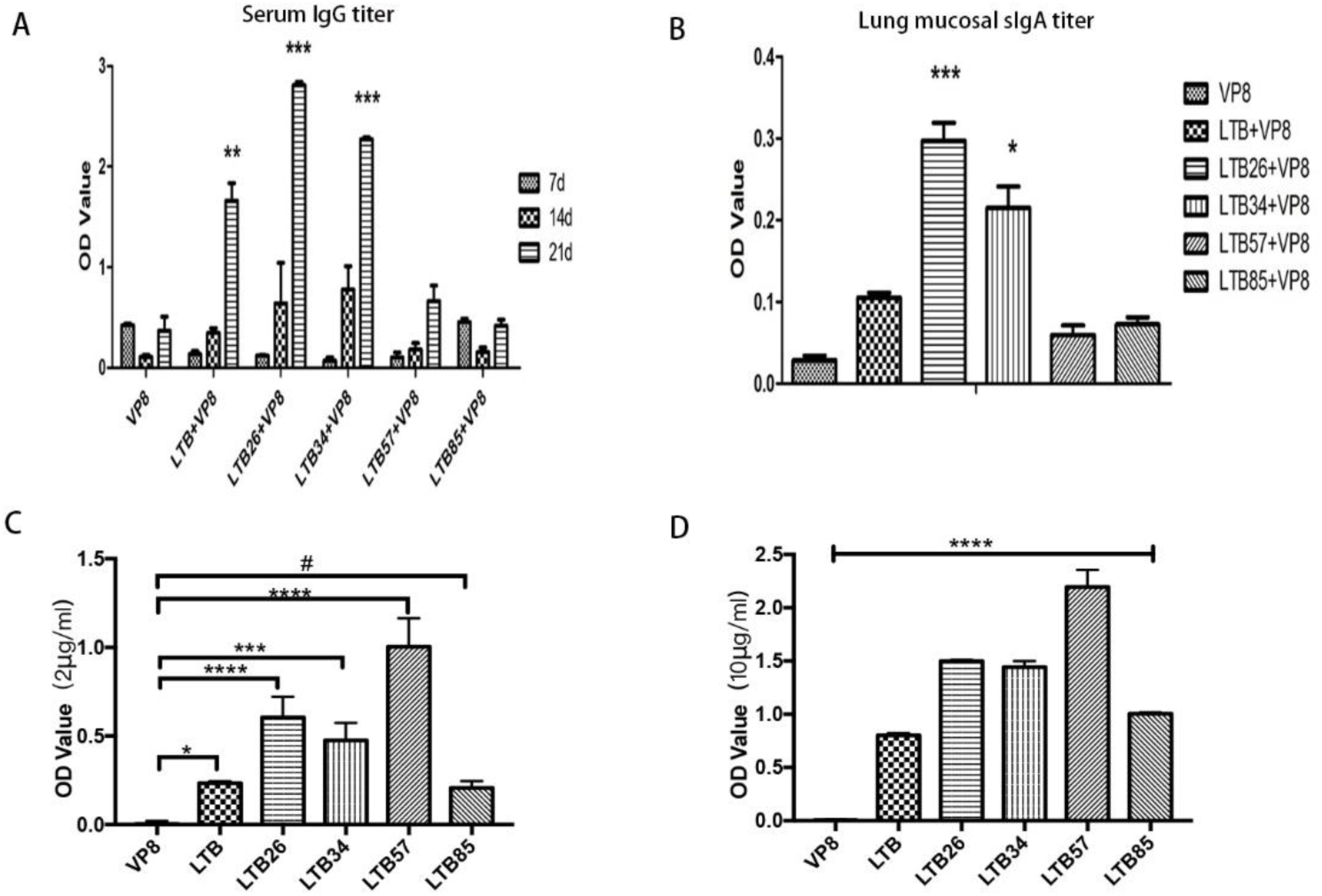
Immune responses and GM1-binding affinity analysis. (A) Serum IgG titer on post-vaccination of 7 d, 14 d and 21 d, respectively; (B) Lung mucosal sIgA titer on the third robust post-vaccination; (C) 1000 ng/well LTB, LTB26, LTB34, LTB57, LTB85, and VP8 were incubated with 2.0 μg/ml GM1. LTB57 had the highest GM1-binding affinity with the lowest adjuvanticity; (D) 1000 ng/well LTB, LTB26, LTB34, LTB57, LTB85, and VP8 were incubated with 10 μg/ml GM1. LTB57 had the highest GM1-binding affinity with the lowest adjuvanticity; ***, p<0.001; *, p<0.01; #, p>0.05.

#### The GM1-binding affinity of LTB mutants no positive association with its adjuvant activity

Previous study suggested that LTB adjuvant positively associated with its GM1-binding affinity(de Haan, Verweij et al., 1998, Holmner et al., 2011). Thus, the GM1-binding activity of the four LTB mutants was tested in turn. The GM1-binding affinity was decreased gradually from LTB57, LTB26, LTB34, and LTB85 to LTB (Fig. 4C, D). However, the adjuvanticity was decreased successively from LTB26, LTB34, LTB, and LTB57 to LTB85. Therefore, LTB26 obtained the highest adjuvanticity with a weaker GM1-binding affinity. Instead, LTB57 had the highest GM1-binding affinity with the lowest adjuvanticity (Fig. 4C,D).

#### The transcriptome analysis indicates LTB26 enhanced B cells via BCR pathway and MHC II^+^ DCs

According to GM1-binding analysis and immune data, purified LTB26 and LTB57 were selected as two models of LTB mutant adjuvants to analyze the differential gene expression. Compared with the group of PBS treated mice, 375 differentiation genes were identified (240 upregulation and 135 downregulation) in the group of LTB+VP8 treatment mice, and 654 differentiations (523 upregulation and 131 downregulation) in the group of LTB26+VP8 treatment. Similarly, 725 differentiations (610 upregulation and 115 downregulation) in the group of LTB57+VP8 treatment, and 890 differentiations (746 upregulation and 144 downregulation) in the group of VP8 alone treatment, respectively (Table 1).

**Table 1.**
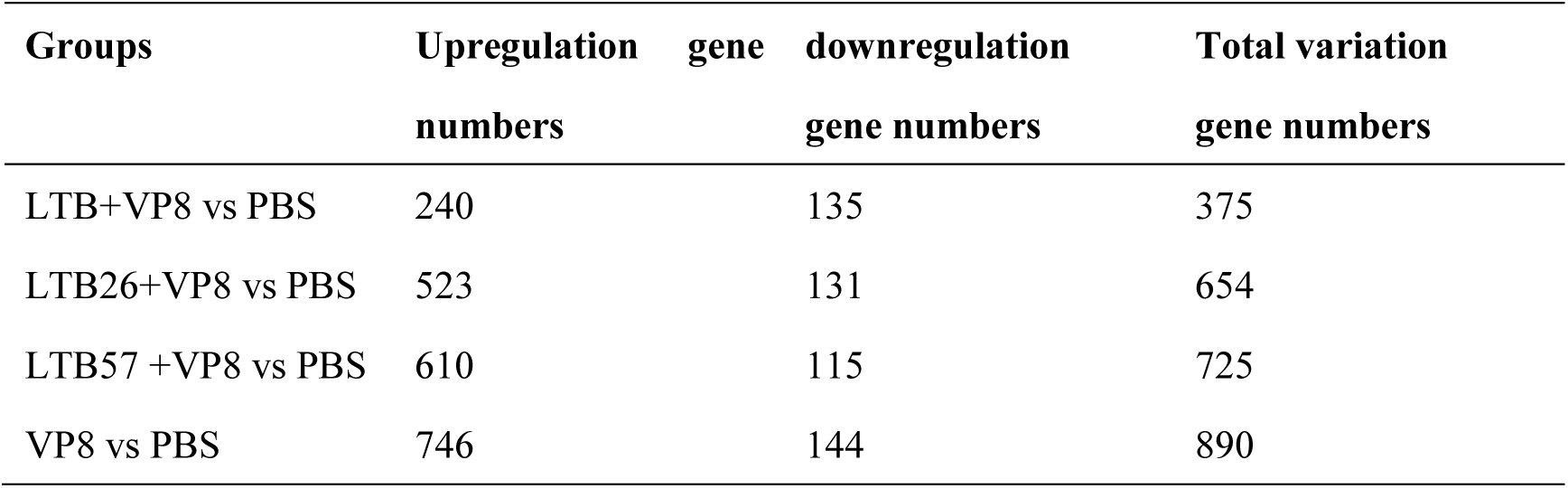
The profiles of differential genes summary.

The Gene ontology (GO) pathway analysis suggested that LTB26 activated several B cell related signal pathways, APC (antigen presenting cell) pathways and MHC II pathway (Fig. 5A). Instead, LTB57 just upregulated inflammatory related signal pathway (Fig. 5B). However, LTB preferred to activate inflammatory response, upregulate TNF and IL-10 expression (Fig. 5C). The results indicated that LTB26 acted as adjuvant via enhancing MHC II related APCs activation and B cell related signal pathway activation (Fig. 5A). That was consistent with the result of SL727-LTB26RBD described above (Fig. 2A, B). Therefore, the molecular mechanism of LTB26 adjuvant was the same, either by binding the receptor into the cell or by expressing it directly in the cell.

**Figure 5.**
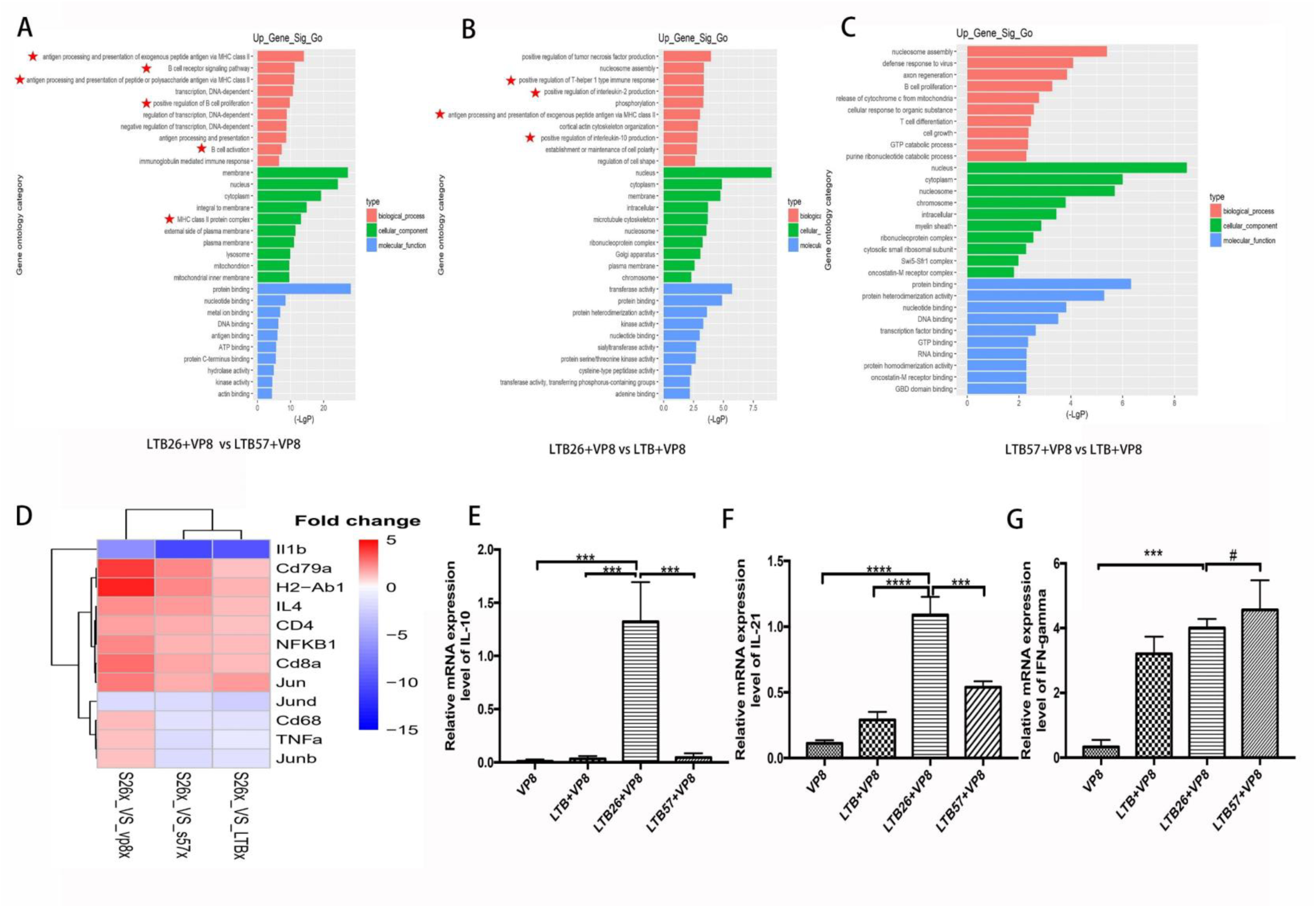
GO analysis the adjuvant related signal pathways and qPCR test. (A)Compared with LTB57, LTB26 significantly activated APCs (MHC II), B cell proliferation, BCR activation and immunoglobulin mediated immune responses, respectively. (B) Compared with LTB, LTB26 significantly activated Th1 immune response, TNF and IL-10 expression, and immune response. (C) LTB 57 did not active APCs (MHC II) and B cell related signal pathways, but upregulated inflammatory response. (D) Cd79a, H2-Ab1, IL-4, CD4, NκkB1, Cd8a, and Jun were significantly upregulated by LTB26. IL-1β and TNFα were significantly downregulated by LTB26. S26: LTB26; S57: LTB57. (E-G) LTB26 significantly upregulated IL-10 and IL-21expression, respectively. LTB26 and LTB57 were significantly upregulated IFN-γ expression than that of VP8, but there were no differences each other. ****, p<0.001; ***, p<0.01; #, p>0.05.

Compared with the expression of immune-related cluster of differentiation (CD) genes, the LTB26 upregulated thirteen B cell associated CDs (CD79a, CD79b, CD19, CD22, CD14, CD37, CD38, CD40, CD48, CD52, CD72, CD74 and CD180), one macrophage marker (CD68) and one common lymphocyte marker (D300c2). Among the fifteen upregulated genes, about 86.7% belonged to B cell associated CDs (Table 2). Of course, LTB upregulated two T cell (and NK cell) marker (CD2, CD8a), one macrophage marker (CD68) and one monocyte, macrophage and DC common marker (CD14). However, B cell marker was no variation in LTB treatment (Table 2). Similarly, LTB57 upregulated three T cell associated CDs (CD47, CD63, and CD164), one lymphocyte homing receptor (CD44), one lymphocyte differentiation marker (CD52), one leukocyte marker (CD53), one neutrophils marker (CD177) and one mast cell marker (D300ld3), respectively (Table 2). The T cell related CDs accounted for about 37.5%. However, LTB57 lost the ability of BCR related activation.

**Table 2.**
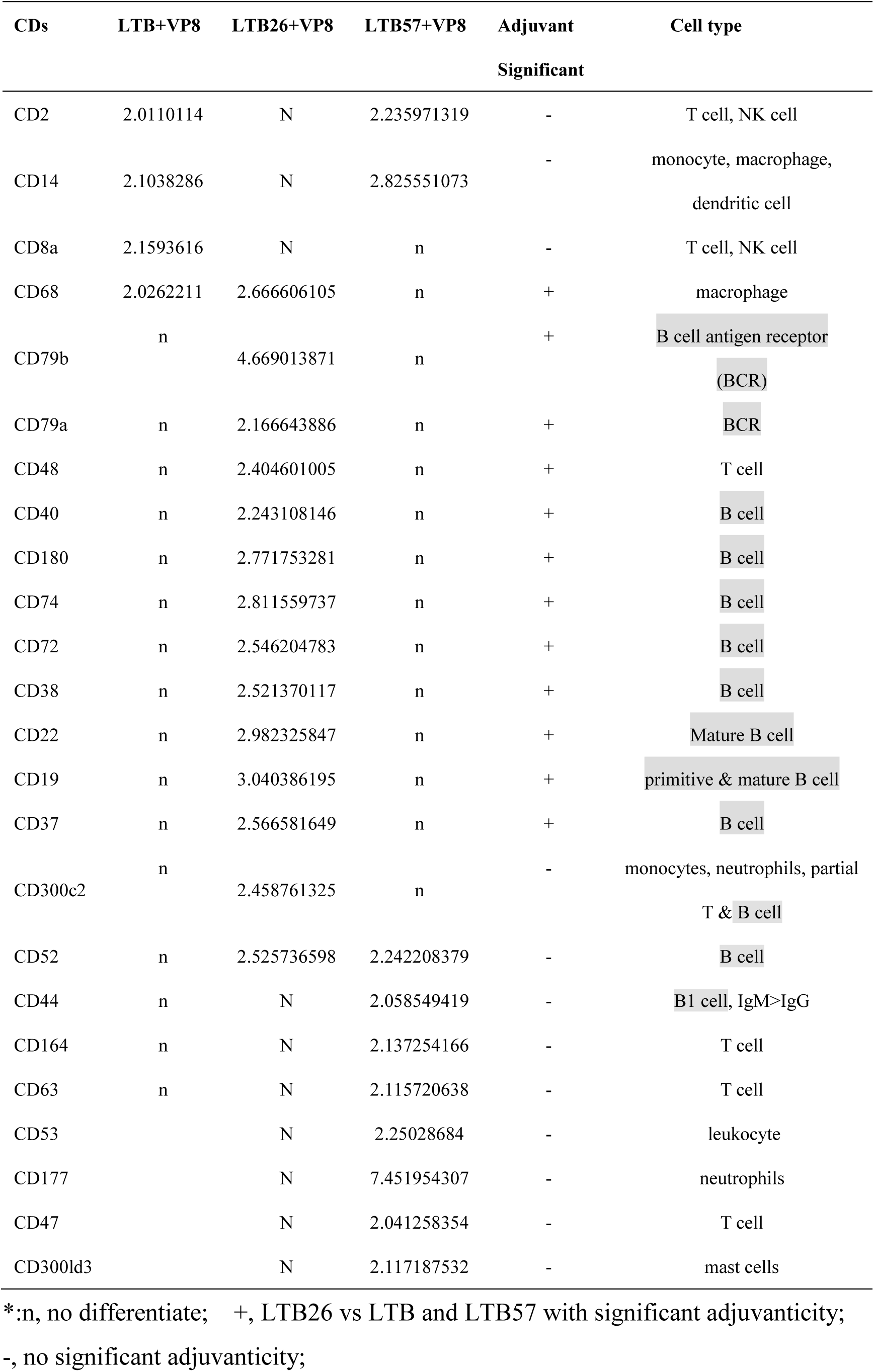
The variation of CDs expression in LTB26 and other groups.

More importantly, CD79a, CD79b and CD19 were the major component of BCR, which indicated that LTB26 functioned adjuvanticity via BCR pathway. The variation of BCR downstream genes showed that LTB26 upregulated the expression of BCR downstream genes more than 2-fold. They were following from Syk, Plc-γ, Ras (Rasa3 and Rras), ERKs (Map3k1 or ERK1), Mapk1ip1l (ERK2), Map4k1 (ERK4)) to transcriptional factors Jund and Atf6b (Table 3). However, LTB57 significantly upregulated the expression of Bcl-10 and the downstream of NFκB inhibitors (Nfκbiz, Nfκbia, and Nfκbib). Otherwise, the upregulation of Mapk6, Akt3 and two transcriptional factors (Egr-1 and Crebrf) did not contribute to adjuvant activity of LTB57 (Supplementary Table 2). Unexpectedly, LTB only upregulated Jun expression (half-level of Jund of LTB26) and significantly enhanced the expression of two NFκB inhibitors (Nfκbia and Nfκbiz) (Table 3). The result suggested that the inhabitation of NFκB activity induced adjuvanticity damage in LTB57 vaccination.

**Table 3.** The variation of BCR downstream genes

| genes | LTB+VP8 | LTB26+VP8 | LTB57+VP8 | Adjuvant<br>Significant |
| --- | --- | --- | --- | --- |
| **Syk** | n | 2.9179075 | 2.8384351 | - |
| **Plcg2** | n | **2.7476057** | n | + |
| Cdc42ep5 | n | 4.1154716 | 7.770395047 | - |
| **Rras** | n | **2.357993** | n | + |
| **Rasa3** | n | **2.2090269** | n | + |
| **Map3k1** | n | **2.115917** | n | + |
| **Mapk1ip1l** | n | **2.1197696** | n | + |
| **Map4k1** | n | **2.2946296** | n | + |
| Mapk6 | n | n | 2.2102702 | - |
| Jun | 2.2490814 | n | n | + |
| **Jund** | n | **5.183886** | n | + |
| Bcl-10 | n | n | 2.0637835 | - |
| Nfkbia | 3.8505929 | n | 5.6544579 | - |
| Nfkbib | n | n | 2.0815464 | - |
| Nfkbiz | 3.3298047 | n | 6.6704407 | - |
| Rasgrp3 | n | n | 2.0318061 | - |
| Atf1 | n | n | 2.1382357 | - |
| **Atf6b** | n | **2.0878157** | n | + |
| Crebrf | n | n | 2.2254203 | - |
| Egr1 | n | n | 3.2136266 | - |
| **Foxo1** | n | **2.5975225** | n | + |
| Akt3 | n | n | 2.4481639 | - |

\*:n, no difference; +, LTB26 vs LTB and LTB26 vs LTB57 with significant adjuvanticity; -, no significant adjuvanticity;

#### Quantitative real-time PCR (q-PCR) verifies the results of transcriptome data

In order to verify the transcriptome data and confirm the above-mentioned APCs, MHC II and BCR pathways, CD8a, CD79a, CD4, IL-1β, IL-4, TNF-α, Jun, Junb, Jund, H2-Ab1, and NFκB1 of PBMC (peripheral blood mononuclear cell) were tested by quantitative real time polymerase chain reaction PCR(q-PCR). The result indicated that ratio of CD4, CD8a, CD79a, IL-4, Jun, H2-Ab1, and NFκB1 were significantly upregulated in LTB26 treatment group compared with LTB and LTB57 treatments, respectively. The increased expression of IL-4 suggested that LTB26 activated Th2 cells response. Meanwhile, the expression of IL-1β, Jund, CD68, TNF-α and Junb were significantly down regulated in LTB26 treatment (Fig. 5D). The result confirmed the transcriptome analysis that LTB26 activated BCR and MHC II pathways and decreased inflammatory response via downregulation of IL-1β and TNF-α.

IL-10 was also commonly produced in Th2 cells and regarded as an anti-inflammatory cytokine and stimulated the proliferation of B cells(Mannino, Zhu et al., 2015, Mocellin, Marincola et al., 2005). Meanwhile, IFN-γ made Naive CD4^+^ T cells differentiate to Th1 cells(Zhang, Zhang et al., 2014). Therefore, IFN-γ, IL-10 and IL-21were also detected by qPCR. Compared with LTB and LTB57 adjuvant, the LTB26 treatment mice were significantly increased expression of IL-10 and IL-21 in lymphocytes (Fig. 3E, F, p<0.001). Because IL-21 played a critical role in T cell-dependent B cell activation(Sondergaard & Skak, 2009, Tangye & Ma, 2020), the result suggested that LTB26 increased the function of T, B and DCs. Similarly, the expression of IFN-γ was significantly increased in LTB26 and LTB57 treatment mice compared with the LTB adjuvant, indicating LTB26 also activated Th1 cells (Fig. 5G, p<0.001). However, there were no significantly change between LTB26 and LTB57 treatment (Fig. 5G, p>0.05). Taken together, LTB26 significantly upregulated both Th1 and Th2 cells activation.

#### The characteristic of FCM of PBMCs

Peripheral blood was sampled to confirm the function of B cell, T cell and APCs elicited by purified LTB and the two mutants (LTB26, LTB57) at 24 h of nasal vaccination. The proportion of CD19^+^CD45R^+^ B cells in BPMCs were 17.82±0.61% (PBS), 24.91 ± 0.82% (LTB26+VP8), 20.08 ± 0.73% (LTB57+VP8), 21.57 ± 1.24% (LTB+VP8) and 20.32 ± 0.61% (VP8), respectively. Compared with VP8 treatment, LTB26+VP8 and LTB+VP8 treatment were significantly increased the proportion of CD19^+^CD45R^+^ B cells in BPMCs than that of LTB57+VP8 vaccination (Fig. 6A, p<0.05, P<0.01), respectively. Meanwhile, there were no significant differences between LTB57+VP8, and VP8 vaccination (Fig. 6A, p>0.05). However, the proportion of CD19^+^CD45R^+^ B cell in groups of LTB26+VP8 and LTB+VP8 were no significantly difference (Fig. 6A, p>0.05).

**Figure 6.**
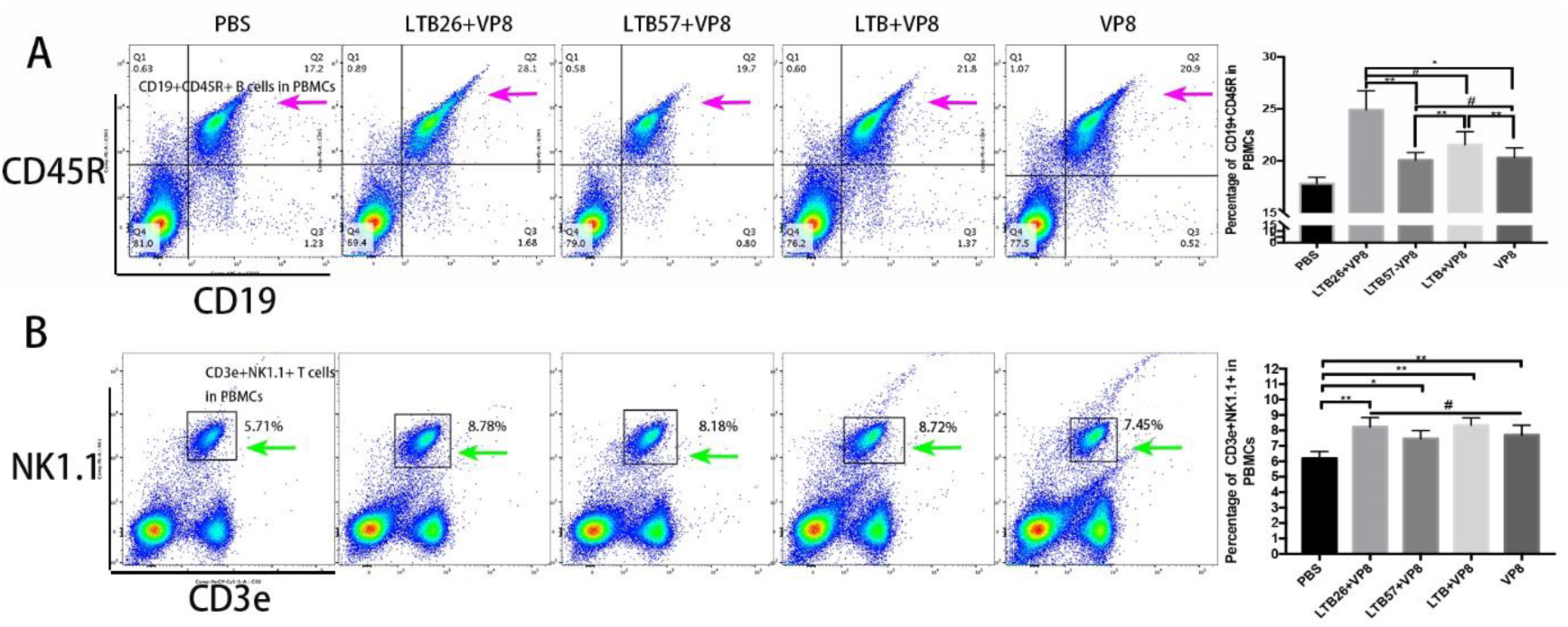
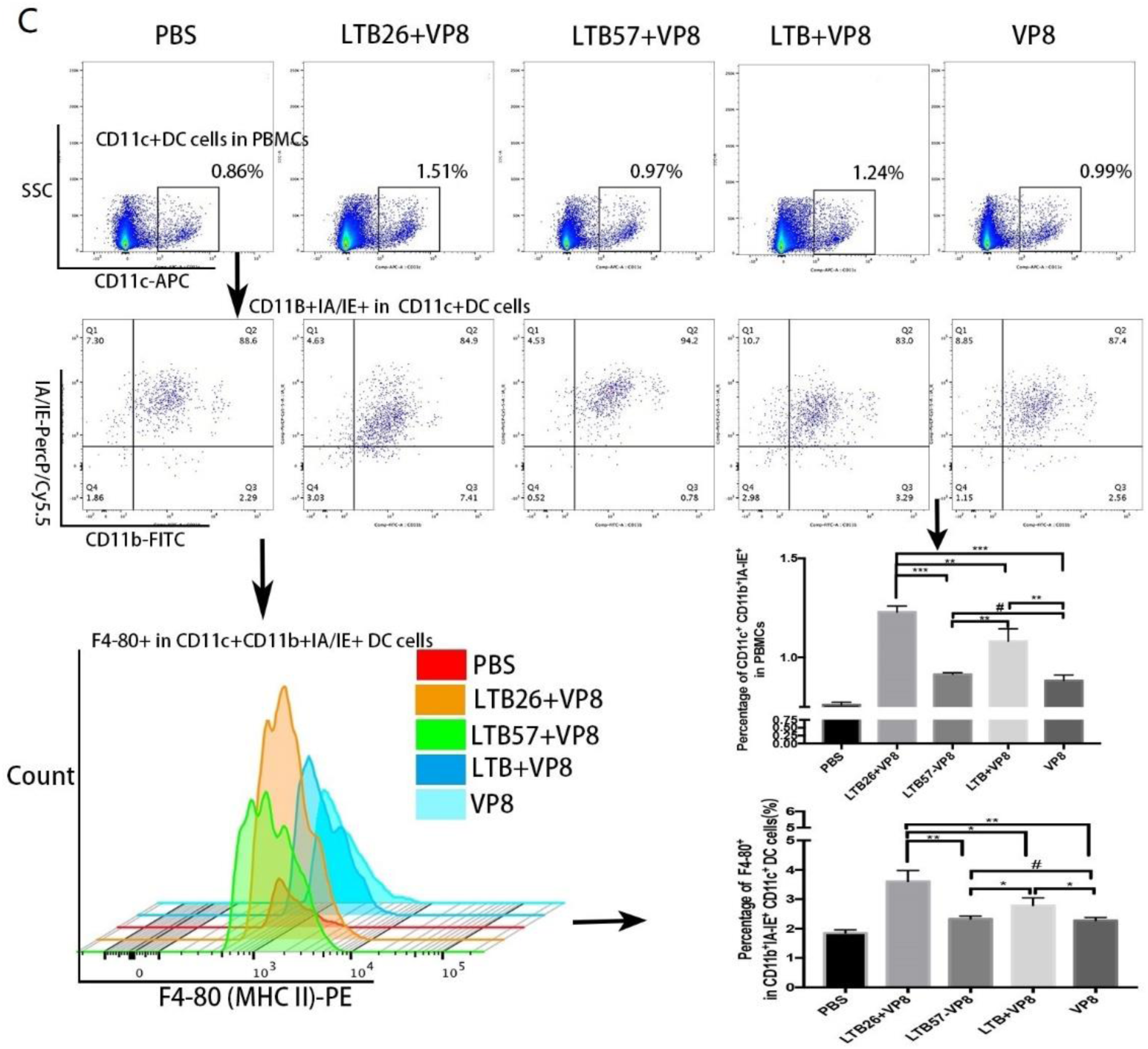
FCM analysis of B cells and NK cells. (A) LTB26 significantly activated CD19^+^ CD45R^+^ B cells. (B) LTB26 significantly activated CD19^+^ CD45R^+^ B cells. (B) NK1.1 cells were no variations among LTB, LTB26, LTB57 and VP8 alone treatment. (C) CD11c^+^CD11b^+^IA/IE^+^ DCs were significantly increased in LTB26+VP8 treatment compared with that of LTB57+VP8, LTB+VP8, and VP8 treatments. **, p<0.001; *, p<0.01; #, p>0.05.

As to the proportion of CD3e^+^ NK1.1^+^ T cells in PBMCs, there were no significant variations among the groups of LTB26+VP8 (8.22±0.60%), LTB57+VP8 (7.47±0.51%), and LTB+VP8 (8.34±0.48%) compared with VP8 treatment (7.71±0.63%), respectively (Fig. 6B, p>0.05).

The proportion of CD11c^+^CD11b^+^IA/IE^+^ DCs in BPMCs were 0.76 ± 0.01% (PBS), 1.23 ± 0.03% (LTB26+VP8), 0.91 ± 0.01% (LTB57+VP8), 1.08 ± 0.06% (LTB+VP8), and 0.88 ± 0.03% (VP8), respectively. The proportion of CD11c^+^CD11b^+^IA/IE^+^ DCs were significantly increased in LTB26+VP8 treatment compared with that of LTB57+VP8, LTB+VP8, and VP8 treatments (Fig. 6C, P<0.001,P<0.01, P<0.001), respectively. That means LTB26 upregulated DCs activation than its wild type.

The F4/80 antigen as a major macrophage marker is expressed on mature macrophages and a subpopulation of DCs (APCs)(Dos Anjos Cassado, 2017). F4/80^+^ cells were major MHC II^+^ mature macrophages with APC function; therefore, the proportion of F4/80^+^ cells in CD11c^+^CD11b^+^IA/IE^+^ DCs was a very important functional indicator of DCs. In this study, the proportion of F4/80^+^ cells in DCs were 1.85±0.11% (PBS),3.61±0.37% (LTB26+VP8),2.33±0.10% (LTB57+VP8),2.79±0.26% (LTB+VP8), 0.28±0.11% (VP8), respectively. Compared with LTB26+VP8 treatment, the proportion of F4/80^+^ cells in DCs were significantly decreased in LTB57+VP8, LTB+VP8, and VP8 treatments (Fig. 6C, P<0.01,P<0.05, P<0.01), respectively. That meant LTB26 were significantly upregulated MHC II^+^ APCs (macrophages and a subpopulation of DCs) activation than its wild type (p<0.05). Summarily, FCM analysis also confirmed the GO data that LTB26 activated MHCII^+^ APCs function and increased B cell activation in line with LTB (wild type).

#### Immunohistochemical staining test confirms the mechanism of LTB26 adjuvanticity

IHC was performed to verify the transcriptome and FCM data. IHC of extent and intensity (EI) score of 0-3 were considered low expression and EI score >3 were considered high expression. The EI values of splenic CD11b were 5.5, 2.875, 2.125, and 1.875 in groups of LTB26+VP8, LTB57+VP8, LTB+VP8, and VP8, respectively. The result suggested that CD11b were significantly upregulated more than 2-fold by LTB26 than that of LTB57, LTB and VP8 treatments (Fig.7A). While the EI values of splenic CD45 were 3, 1.125, 1.25, and 1.375 in groups of LTB26+VP8, LTB57+VP8, LTB+VP8, and VP8, respectively. The result suggested that CD45 were significantly upregulated more than 2-fold by LTB26 than that of LTB57, LTB and VP8 treatments (Fig. 7A). Of course, the level of CD45 was slightly decreased in LTB26+VP8 treatment than that of CD11b. The EI values of splenic CD4 were 4.125, 3.75, 1.625, and 1.875 in groups of LTB26+VP8, LTB57+VP8, LTB+VP8, and VP8, respectively. The result suggested that CD4 were significantly upregulated more than 2-fold by LTB26 and LTB57 than that of LTB treatments (Fig. 7A). Nonetheless, the EI value of LTB26 treatment was higher than that of LTB57.

**Figure 7.**
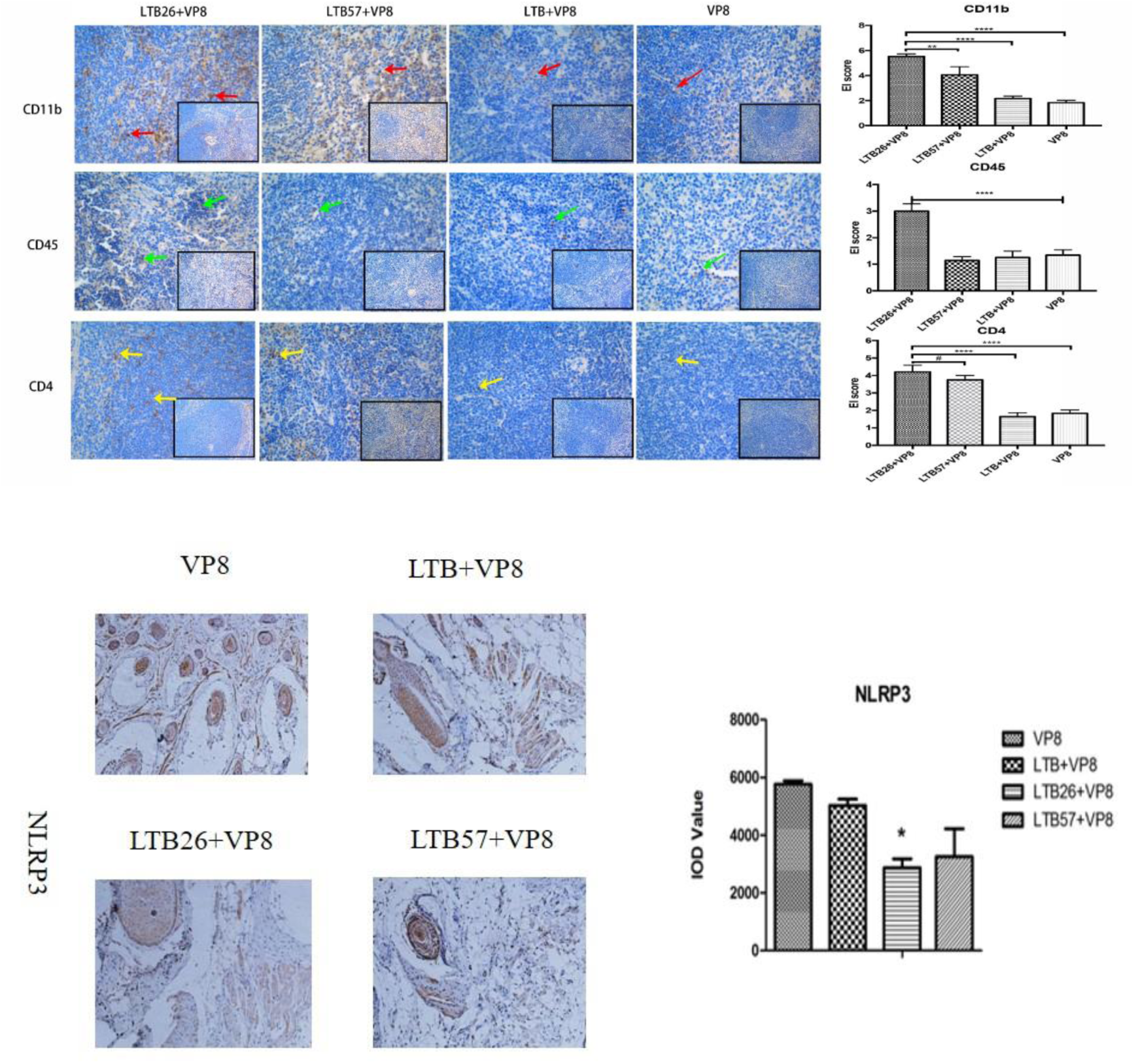
IHC analysis of CDs and NLRP3. (A) CD11b, CD45, CD4 were significantly upregulated more than 2-fold by LTB26. (B) NLRP3 was significantly downregulated by LTB26.****, p<0.001, **, p<0.01; #, p>0.05.

NLRP3 was one of NOD-like Receptors (NLRs) and played a crucial role in Alum-based adjuvant immune by activating of NLRP3 inflammasome and IL-1β production via MAPK signaling pathway(Sun, Ji et al., 2017). However, the expression of NLRP3 in nasal tissue of the purified LTB, LTB26 and LTB57 adjuvant mice was lower than that of in VP8 alone treatment (Fig. 7B, p<0.01). The result indicated that LTB and its mutants functioned mucosal adjuvant independent on NLRs activation.

### Modeling the three-dimensional structure of LTB26

The wild type of LTB hit a protein named subunit of heat-labile enterotoxin B (template serial number: 5lzi.1.A) as a reference template for this modeling in SWISS-MODEL database. Then 3D model of LTB26, LTB34, LTB57 and LTB85 monomer and pentamer structures were established (Fig. 8 A, B, C, D, E, F, G, H, I, J). The result showed that LTB26 changed the first α-helix of LTB (epitope P1, EP1) to a β–sheet and endowed LTB26 more higher adjuvanticity than LTB with unknown mechanisms (Fig. 3D)(Guan, Liu et al., 2015, Takahashi et al., 1996). Intriguingly, the octapetide (M26S27N28T32I33N34) mutation of LTB26 enhanced the GM1-binding capacity instead. It indicated that the first α-helix of LTB hindered its adjuvanticity, not its amino acid sequences. LTB34 mutation covered three amino acid of the C-terminal of EP1 and three amino acid of the N-terminal of EP2. However, the decapeptide mutation (L35S36L37S38N39S40T41I42N43Y44) of LTB34 were maintained the structure of the second β–sheet and impaired neither GM1-binding affinity nor adjuvanticity (Fig. 3D). Thus, this fragment was no contribution to both of GM1-binding affinity and adjuvanticity of LTB. The residue of Y39 of LTB was reported to involve in stabilizing the GM1-binding(Holmner et al., 2011, Ma, 2016). It was suggested that Y39N mutation did not damage it stabilizing effect.

**Figure 8.**
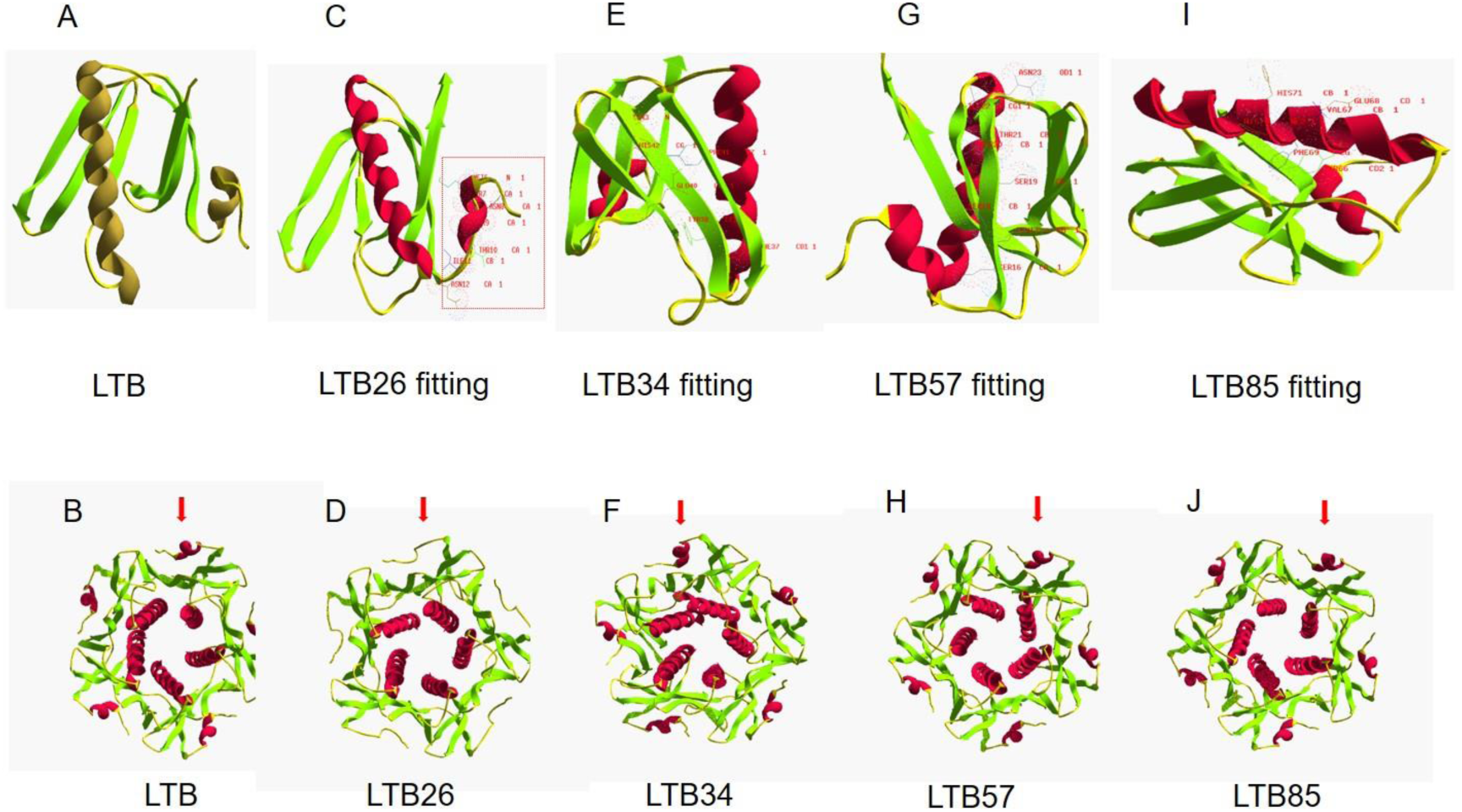
Prediction of LTB and LTB mutants 3D structure. (A) 3D structure of LTB monomer. (B) 3D structure of LTB pentamer. (C) 3D structure of LTB26 and LTB fitting monomer. The first α-helix of LTB was replaced with β-sheet in LTB26 (red box) (D) 3D structure of LTB26 pentamer. Arrow showed the β-sheet. (E) 3D structure of LTB34 and LTB fitting monomer. The 3D structure was intact. (F) 3D structure of LTB34 pentamer. Arrow showed the first α-helix. (G) 3D structure of LTB57 and LTB fitting monomer. The 3D structure was intact. (H) 3D structure of LTB57 pentamer. Arrow showed the first α-helix. (I) 3D structure of LTB85 and LTB fitting monomer. The 3D structure was intact. (J) 3D structure of LTB85 pentamer. Arrow showed the first α-helix.

Instead, the LTB57 mutant maintained the structure of the third β–sheet similar to LTB, however, the amino acids mutation led to lose its adjuvanticity. This mutation (F57Y58E60F61H62H63) was located in the central of epitope EP3 (48-63) and EP4 (54-69) and significantly impaired the adjuvanticity (Fig. 3D)(Guan et al., 2015). However, it was separated two residues from the downstream of the typical GM1-binding residue of G33 (numbered G54 in this study) (Fig.3B, C, D)(Ma, 2016, Nashar, Webb et al., 1996, Turcanu, Hirst et al., 2002). Similarly, LTB85 (86-91 aa) mutation was also located in the central T- and B-cell epitope EP5 and retained the second α-helix of LTB(Guan et al., 2015, Ma, 2016). Even though the K91 of LTB57 (K91H) was a conserved GM1-binding residue, the hexapetide mutation (Y86V87E88F89H90H91) was not impaired the GM1 affinity, too (Holmner et al., 2011, Ma, 2016). However, both LTB57 and LTB85 lost their adjuvanticity for destroying the sequences of T- and B-cell epitope of LTB (Fig.3C, D). Therefore, the sequences of epitopes of EP3, EP4 and EP5 were important than that of 3D structures for maintaining the adjuvanticity of LTB.

## Discussion

COVID-19 was the most dangerous infectious disease to threat public health since December 2019 causing more than 89 million cases infection with more than 2.1 million death to date(Harrison, Lin et al., 2020). Vaccines could help prevent the spread of COVID-19 and end this pandemic. Since on Dec. 18, 2020, FDA had approved two COVID-19 vaccines, several vaccines have been approved and vaccinated in many countries. However, the constant mutation of SARS-COV-2 was not only challenged the effectiveness of the vaccine, but also increased its infectivity(Chen, Wang et al., 2020, Plante, Liu et al., 2020). Previous study suggested that G614 mutation might reduce the ability of vaccines in clinical trials to protect against COVID-19(Plante et al., 2020). Unfortunately, several variations (A348T, R408I, K417N, G476S, V483A, E484K, N501Y, H519Q, A520S) were found at the RBD region (328-533aa, DOI:10.1101/2020.12.24.20248822v1; DOI: 10.1101/2020.12.21. 20248640)(Gu & Chen, 2020, Lokman, Rasheduzzaman et al., 2020). N501Y mutant was recently reported with the increased virulence and pandemic worldwide(Gu & Chen, 2020). A strain, VOC 202012/01, was found in British accumulated 17 amino acid mutations including 69-70del, N501Y, P681H, and other 4 mutations in spike protein (DOI:10.1101/2020.12.24.20248822v1). The 69-70 deletion was linked to immune escape in immunocompromised patients and enhanced viral infectivity in vitro (DOI:10.1101/2020.12.24.20248822v1). Other mutants with K417N or E484K were found in South Africa.

Therefore, it remained to be monitored whether the current vaccines would be effective against these mutants in the future. Therefore, the battle against COVID-19 would continue for some time because of insufficient supplies of vaccines. However, RBD was still the effective antigen for vaccine development. We, based on this, developed an attenuated *Salmonella typhimurium*-based RBD oral vaccine fusing LTB26 as mucosal adjuvant. Then the RBD mutants’ vaccine will be constructed on the basis of it in the future.

*Salmonella ssp* was broadly used as oral vaccine delivery(Yurina, 2018). Attenuated *Salmonella typhimurium* was the most widely used vectors. It was suitable for oral immunization and boosted both mucosal and systemic immune responses(Yurina, 2018). Therefore, as a mucosal infectious pathogen, *Salmonella*-based oral vaccine was suitable for SARS-COV-2 prevention. Another advantage of *Salmonella*-based oral vaccine was that the recombination delivery vehicle was easy to construct. Meanwhile, *Salmonella*-based oral vaccine was essentially DNA vaccine and the antigenicity of RBD was same as the production of infectious SARS-COV-2 spike protein. Although *Salmonella*-derived PAMPs (pathogen-associated molecular patterns) has adjuvant activity, we found that its adjuvanticity was weaker than LTB26 (Fig. 2)(Bivona, Cerny et al., 2016). Therefore, we designed LTB26RBD fusion gene to enhance the immune response in this study.

Mucosal immunity was one of the important ways to protect the body from the mucosal invading pathogens(Lycke, 2012, Ma, 2016, Perez-Lopez, Behnsen et al., 2016). However, mucosal adjuvant research lagged behind vaccine development and resulted in vaccines failure in some case(Atmar & Keitel, 2009, Young, Sadarangani et al., 2015). Therefore, it was also important to develop safe and high effective mucosal adjuvants to enhance the effectiveness of mucosal vaccines.

LT could regulate the differentiation of B cells and regulate the production of T cells through mucosal immunity(da Hora et al., 2011, Holmner et al., 2011). However, the toxicity of LT prevented its application as adjuvant in human. In recent years, non-toxic LTB adjuvant had been widely used(Cunha, Moreira et al., 2017, Ghazali-Bina et al., 2019, Holmner et al., 2011, Liu, Ma et al., 2016, Ma, Luo et al., 2012). The binding specificity of LTB to GM1 was associated with adjuvant activity. Previous studies reported that the property of LTB adjuvant was dependent on GM1- binding affinity(de Haan et al., 1998). The LTB (G33D, numbered G54 in this study for counting the N-terminal signal peptide) mutant might lose the immune adjuvant activity for failing bind to GM1(de Haan et al., 1998). However, GM1 binding sites also existed in other epitopes of LTB such as 51E, Ile58, 58I and 91K(Holmner et al., 2011). Therefore, we had mentioned that beyond GM1-binding, G33 of LTB was a crucial antigenic determinant residue located in the B- and T-cell epitope region (residue S26 to G45)(Ma, 2016). Scientist supposed that G33 residue was crucial for the binding affinity of LTB to GM1 and LTB (G33D) mutant lost its GM1-binding affinity and oral immune adjuvant activity(Nashar et al., 1996).

Even so, we had deduced that LTB (G33D) destroyed the function of the key residue of LTB antigenic determinant (residue S26 to G45) rather than the GM1-binding affinity(Ma, 2016). Therefore, the integrity of LTB epitope was more important than the GM1-binding affinity(Ma, 2016). This hypothesis has been verified that LTB57 (57-63 aa) mutation (F57Y58E60F61H62H63) was located in the central T- and B-cell epitope and significantly impaired the adjuvanticity (Fig.3D)(Ma, 2016, Nashar et al., 1996, Turcanu et al., 2002). Intriguingly, the octapetide (M26S27 N28--T32I33N34) mutation in LTB26 was not only impaired the GM1 affinity, but also enhanced the GM1-binding capacity. Similarly, LTB85 (86-91 aa) mutation was also located in the central T- and B-cell epitope(Ma, 2016), and this hexapetide (Y86V87E88F89H90H91) mutation was not impaired the GM1 affinity, too. Even though the K91 of LTB57 (K91H) was a conserved GM1-binding residue(Holmner et al., 2011, Ma, 2016). However, both LTB57 and LTB85 lost their adjuvanticity for destroying the structure of T- and B-cell epitope of LTB (Fig.3D). Therefore, the highlight of this work suggested that the adjuvanticity of LTB was not positively associated with GM1-binding affinity.

Adjuvant stimulated APCs and targeted the innate immune system to induce a robust immune response. In principle, adjuvant could induce host-derived damage-associated molecular patterns (DAMPs) or recognizing PAMPs to enhance antigen-specific immune responses via APCs (DCs) or macrophages^(Hayashi, Momota et al., 2018)^. DCs sensed and phagocytized invading pathogens and activated naïve T cells. Then they acted as a major link between innate and acquired immunity to determine the polarization of T cell responses into different effector subtypes(Tesfaye, Gudjonsson et al., 2019) (Steinman & Hemmi, 2006).

Cytokines were important immune regulators. Purified LTB26 was capable of orchestrating upregulated IL-4, IL-10 and IL-21 expression, significantly, compared with LTB and LTB57 (Fig. 5D, 5E, 5F). Generally, B cells and CD8^+^ T cells were the responders to IL-21 by IL- 21R(Sondergaard & Skak, 2009). CD4^+^ T cells were main producers of IL- 21, while TCR stimulation increased CD4^+^ T cell expression of IL-21R as a positive feedback. IL-10 stimulated cytotoxicity of CD8^+^ T cells and IFN-γ expression(Oft, 2014).

IL-4 promoted humoral immunity and IFN-γ promoted cell-mediated immunity(El-Sissi, Mohamed et al., 2020). LTB, LTB26 and LTB57 significantly upregulated the expression of IFN-γ and TNF-α compared with VP8 alone treatment, but there were no differentiation among LTB, LTB26 and LTB57 (Fig. 5G). Thus, purified LTB26 promoted both Th1 and Th2 cells mediated immunity via upregulating IL-4 and IFN-γ expression, which was different from Alum-based vaccine (IL-4 only). This consistent with the data obtained from FCM and transcriptome analysis.

Together our data indicated that LTB26 adjuvant fused with SARS-COV-2 RBD or purified and mixed with hRV VP8 significantly increased the proportion of MHC II^+^ DCs and CD19^+^CD45^+^ B cells in peripheral blood, which indicated that LTB26 enhanced the processing of vaccine (antigen) via DCs activation and the producing of antibody by B cell activation (Fig 1, 6). We found *Salmonella typhimurium* SL7207 was not an ideal vehicle for oral vaccine development for its toxicity. In this study, the BALB/c mice and SD rats were died within 24 h after intraperitoneally injection with the same dose of oral vaccination SL7207 (data not show). Therefore, the less toxicity *Salmonella* strain or other vehicles were necessary in future research. Furthermore, the mucosal responses of SL7207-LTB26RBD will be performed as a single topic in the future.

## Material and Methods

### Construction of pcDNA3.1-LTB26RBD plasmid

LTB26 was derived from LTB (Genbank No. EU113252) with I26MT27SE28NL29ΔC30Δ (L29C30 deletion) and E32TY33IH34N mutation. SARS-COV-2 RBD (301-521 aa, Genbank No. YP_009724390) was fused with GSGSG linker at C-terminal of LTB26. The sequence of LTB26RBD and RBD were commercially synthesized and inserted into pcDNA3.1(+) plasmid at *Bam*H I/*Not* I position to construct pcDNA3.1-LTB26RBD and pcDNA3.1-RBD plasmids, respectively. Then the plasmids were electrically transformed into attenuated *Salmonella typhimurium* SL7207 (SL7207) as oral vaccine candidates (SL7207-RBD, SL7207-LTB26RBD) at 2.5kv and confirmed by sequencing.

### Animal and Immunization

BALB/c mice of 3-4 weeks (male) were bred in the experimental animal center of Chongqing Medical University and divided into four groups. Six groups of mice (n=5) were oral vaccinated with *Salmonella* SL7207, SL7207-RBD, SL7207-LTB26RBD and 200 μl PBS (2.5×10^6^ cfu /mouse), respectively, after treating with saturated sodium bicarbonate (0.2 ml/mouse). The mice were boosted twice on 10, and 20 day as described above. Blood was sampled from the orbital vein on day 0, 10, 20, 30.

This study was carried out in strict accordance with the recommendations in the Guide for the Care and Use of Laboratory Animals of the National Institutes of Health. The protocol was approved by the Committee on the Ethics of Animal Experiments at the Chongqing Medical University (SYXK2012-0001, 2013-03-11).

### Flow Cytometry

PBMCs from heparinized blood were isolated with Ficoll-Paque. The PBMC samples were stained with 5 µl of anti-mouse CD3e-PerCP/cy5.5 (Biolegend, USA, cat#100327), NK1.1-PE (Biolegend, USA, cat#108707),CD19-PE (Biolegend, USA, cat#115511), CD45R/B220-APC (Biolegend, USA, cat#103207), CD11c-APC (Biolegend, USA, cat#117309), CD11b-FITC (Biolegend, USA, cat# 101205), F4/80-PE (Biolegend, USA, cat#123109), and IA/IE-PerCP/cy5.5 (Biolegend, USA, cat#107625), respectively, for 30 min at RT in the dark. The cell were stained with 5 µl of PerCP/cy5.5-Armenian Hamster-IgG (Biolegend, USA, cat#400931), PE-Rat-IgG2a (Biolegend, USA, cat#400508), PE-Rat-IgG2b (Biolegend, USA, cat#400211), APC-Rat-IgG2a (Biolegend, USA, cat#400511), APC-Rat-IgG1 (Biolegend, USA, cat#401903), FITC-Rat-IgG2b (Biolegend, USA, cat#400634), PerCP/cy5.5-Rat IgG2b (Biolegend, USA, cat#400631), and respective isotype control for 30 min at RT in the dark, respectively. Then washed twice with 500 µl PBS and assessed by four-colored flow cytometry. Then measured the percentages of CD3e^+^NK1.1^+^ NK cells, CD19^+^CD45R^+^ B cells and CD11c^+^CD11b^+^F40^-^80^+^IA/IE^+^ DCs (dendritic cells) and the fluorescence intensity (MFI) of cell finally.

### SL7207-LTB26RBD immune assay

Blood samples were individually collected from oral immunized mice by orbital venous bleeding on days 0, 10, 20 and 30 for the analysis of systemic RBD specific antibodies (n=5). All the samples were treated as previously described(Liu et al., 2016). The antibodies were analyzed with horseradish peroxidase (HRP)-labeled goat anti-mice IgG and goat anti-mice IgA (1.0 μg/ml, Boster, Wuhan, China) by ELISA, respectively, as previously described(Liu et al., 2016). SARS-CoV-2 RBD-His protein was purchased from Nanjing Okay Biotechnology Co., Ltd (Nanjing, China, cat#K1516). Endpoint titers were determined as the dilution of each sample showing a 2.1-fold higher absorbance level of 450 nm as compared to that of the negative control samples. Average OD_450_ values for the animals were calculated.

### LTB mutants design

Full-length LTB DNA was cloned from EC44815 strain and constructed pET32a-LTB plasmid as our previously report(Ma et al., 2012). The four LTB mutants were designed to mutate some amino acids located in the B- and T-cell epitope region(Ma, 2016). The amino acid was numbered from the first M (Met) of signal peptide in LTB. The four LTB mutants were also constructed into plasmid pET32 at *Bam*H I /*Sal* I site, respectively. Full-length of hRV VP8 DNA (GenBank: L34161) was commercially synthesized (Sangon, Shanghai, China) and inserted into plasmid pET32 at *Bam*H I/*Sal* I site. The recombinants of pET32-LTB, pET32-LTB26, pET32-LTB34, pET32-LTB57, pET32-LTB85 and pET32-VP8 were expressed in *E. coli* BL21 cells and purified with BeaverBeads™ His-tag protein purification kits (Beaver, Suzhou, China, cat#70501-5). The endotoxin was removed using ToxinEraserTM resin (Genscript, Nanjing, China, cat#L00338) and the protein concentration was measured by BCA protein assay kit (TaKaRa, Dalian, China, cat#T9300A) as previous described(Ma et al., 2012).

### Animal and Immunization

Six groups of BALB/c mice (3-4 weeks old, male, n=6) were nasal vaccinated with human rotavirus VP8 (VP8), LTB+VP8, LTB26+VP8, LTB34+VP8, LTB57+VP8, LTB85+VP8, and 20 μl PBS, respectively, after anesthetizing with chloral hydrate (0.5 mL/100 g). The VP8, LTB and four LTB mutants were each added 10 μg/mouse respectively. The mice were intranasal boosted twice on 7, and 14 day with same method after first vaccination. All surgery was performed under sodium pentobarbital anesthesia, and euthanized by cervical dislocation. Then immune assay was performed as described above to detect the VP8 specific antibodies from blood, fecal and bronchial mucosal washing, respectively. The PBMCs FCM analysis was performed as previous described.

### GM1-binding analysis

The GM1-binding affinity of LTB and its four mutants were determined using GM1-ELLSA (enzyme-linked immunosorbent assay) assay^(Minke, Roach et al., 1999)^. The wells of microplate were coated with 200 µL (2.0 µg/ml and 10 µg/ml, respectively) of GM1 (Qilu pharma, Shandong, China) or PBS, respectively, at 48℃ overnight. Plates were washed three times with 500 µl PBS to remove uncombined GM1. Subsequently, plates were blocked by the addition of 200 µl PBS containing 1.0 % BSA at 37℃ for 30 min. Then, plates were washed again as described above. Finally, plates were incubated with 1000 ng/well LTB, LTB26, LTB34, LTB57, LTB85, and PBS, respectively, at 37℃ for 2 h, followed by washing as described above. 100 µl of rabbit anti-His-tag antibody (1:1000, BioVision, USA, cat#3998) was incubated at 37℃ for 2 h and washed as described above. 100 µl of HRP conjugated goat anti-mouse IgG(H+L) secondary antibody (1:1000, Boster, China, cat# BA1051) was incubated at 37℃ for 2.5 h, followed by washing four times. Finally, added 100 µl of TMB (tetramethylbenzidine) to each well and incubated at 37℃ for 5 min. Then the reaction was stopped by 200 µl H_2_SO_4_ (2.0 mol/l). The OD_450_ was read by microplate Reader.

### Transcriptome analysis of peripheral lymphocyte

According to animal immune data, the peripheral lymphocytes were sampled in groups of LTB, LTB26+VP8 (the highest immune adjuvanticity with middle GM1-binding capacity), LTB57+VP8 (the highest GM1-binding affinity with lowest immune adjuvanticity), VP8 (in PBS), and PBS treatment mice after 24 hours vaccination. Total RNA was extracted using TRIzol Reagent (Invitrogen, USA) according the manufacturer manual. The RNA quality were measured using a NanoDrop 2000 spectrophotometer (Thermo Fisher, USA) at 260/280 nm and agarose gel electrophoresis. Then RNA quantity was measured using Qubit2.0 Fluorometer. mRNA sequencing was performed on Illumina Hiseq platform (Illumina, USA) with 12 G bps and 10 M reads (Genminix Informatic Ltd., Shanghai, China). The differentially expressed genes were selected as having more than 2- fold difference between their geometrical mean expression in the compared groups and a statistically significant P-value (<0.05) by analysis of DEseq2. The GO analysis on differentially expressed genes was performed with an R package: cluster profiler using a P<0.05 to define statistically enriched GO categories. Pathway analysis was used to determine the significant pathway of the differential genes according to Kyoto Encyclopedia of Genes and Genomes Database (http://www.genome.jp/kegg/).

### Peripheral blood qPCR analysis

Total RNA was isolated as previous described. RNA reverse transcription was performed using the TaqMan RNA Reverse Transcription kit (Thermo Fisher, USA, cat#4366596). qPCR was performed using SYBR Green Supermix (Bio-Rad, USA, cat# 1708882) and the iCycler iQ Real-Time PCR system (Bio-Rad, USA). The reactions were run as follow: denaturation at 95°C for 10 min followed by 50 cycles of 95°C for 10 sec, 55°C for 10 sec and 72°C for 5 sec; 99°C for 1 sec; 59°C for 15 sec; 95°C for 1 sec; then cooling to 4°C. Relative mRNA expression was normalized against the endogenous control, GAPDH, using the 2-ΔΔCt method. The primers used in the current study were listed as table (Supplementary Table 1).

### Immunohistochemistry

Spleen played an important role for lymphocyte migration and immune response after receiving antigen stimulation. Therefore, CD4, CD11b, and CD45 of splenic lymphocyte were tested after 24h poster vaccination. Haematoxylin and eosin staining and antibody labeling were performed on 4-μm tissue sections as described previously(Liu et al., 2016). Antibodies(rabbit) against CD11b, CD45 and CD4 were incubated overnight at 4 ℃. Antibodies (goat) against IgG (rabbit) were used for labeling for 2 h at room temperature. Images of histology and immunohistochemistry were taken with a Nikon Eclipse E600 and Nikon DS-Ri1 camera or with a Digital Slide Scanner (20 × magnification).

The expressions of CD11b, CD45 and CD4 were quantified using a visual grading system based on the extent of staining as previously described(Marechal, Demetter et al., 2009). Briefly, percentage of positive spleen cells (extent of staining) was graded on scale from 0 to 3: 0, none; 1, 1-30%; 2, 31-60%; 3, 41-60%. The intensity of staining was graded on a scale of 0-3: 0, none; 1, weak staining; 2, moderate staining; 3, strong staining. The combination of extent (E) and intensity (I) of staining was obtained by the product of E times I called EI varying from 0 to 9 for each spot. For statistical analysis, EI score of 0-3 were considered low expression and EI score >3 were considered high expression.

### Three-dimensional structure prediction

In order to understand the enhanced adjuvanticity of LTB26, the three-dimensional structures of LTB26, LTB34, LTB57 LTB85 and LTB were analyzed using SWISS-MODEL project (https://swissmodel.expasy.org/interactive/HTEqUf/models/). The quality of model was evaluated using the SWISS-MODEL scoring function of GMQE, QMEAN4, Identity, and Method parameters. GMQE [0, 1] score estimation results of the tertiary structure of the model accuracy, the higher the better; QMN4 is to evaluate the global and local quality of the model to ensure the reliability of the overall quality of the model. QSQE combines interface protection, structural clustering, and other template features to provide quaternary structural quality estimates that reflect the expected accuracy of interchain contact based on a given alignment and templated model. The higher the number, the higher the reliability, which complements the GMQE score. Identity stands for sequence alignment matching rate, the higher the better.

## Acknowledgements

The work was funded by a research grant from the National Natural Science Foundation of China (NSFC no. 30972585); We thank professor Shengyong Yang (Chongqing Medical University) proofread the manuscript and gave some constructive advises.

## Contributions

Y.M. conceived of the paper; Q.S. and Q.W. contributed equally and performed main experiments; S.C., S.G. and C.F. performed some experiments; Y.M., Q.S., F.S. and T.L. contributed to write the paper.

**Supplementary material:** Supplementary Table 1. qPCR primers used in this study.

