## supplemental Table 1, 2 for "SARS-COV-2 RBD Oral Vaccine Boosted by Mucosal Immune Adjuvant LTB26 via DCs and B Cells Activation in Mice"

**Supplementary Table 1. The profiles of differential genes summary**

| Groups | Upregulation gene numbers | downregulation gene numbers | Total variation  gene numbers |
| --- | --- | --- | --- |
| LTB+VP8 vs PBS | 240 | 135 | 375 |
| LTB26+VP8 vs PBS | 523 | 131 | 654 |
| LTB57 +VP8 vs PBS | 610 | 115 | 725 |
| VP8 vs PBS | 746 | 144 | 890 |

**Supplementary Table 2. Primers for qPCR**

| **Genes** | **Primer sequences** |
| --- | --- |
| Cd8a | Forward: 5’-tcagttctgtcgtgccagtc-3’  Reverse: 5’-atcacaggcgaagtccaatc-3’ |
| Cd68 | Forward: 5’- tagcccaaggaacagaggaa-3’  Reverse: 5’- ggagctggtgtgaactgtga-3’ |
| Cd79a | Forward: 5’- ggtaccaagaaccgcatcat-3’  Reverse: 5’- caaggttcaggccctcatag -3’ |
| CD4 | Forward: 5’- aggaagtgaacctggtggtg- 3’  Reverse: 5’- ctcctgcttcagggtcagtc -3’ |
| Il-1b | Forward: 5’- gaccttccaggatgaggaca -3’  Reverse: 5’- agctcatatgggtccgacag -3’ |
| IL-4 | Forward: 5’- cctcacagcaacgaagaaca -3’  Reverse: 5’- ctgcagctccatgagaacac -3’ |
| IL-10 | Forward: 5’-tccttggaaaacctcgtttg-3’  Reverse: 5’-cttcaattgcttcccaagga-3’ |
| IL-21 | Forward: 5’-attaaagcttctggtggcatggagaggac-3’  Reverse: 5’-taggatcctgtgttctaggagagatgctg-3’ |
| IFN-γ | Forward: 5’-gtgattgcggggttgtatct -3  Reverse: 5’-tgtcattcgggtgtagtcaca -3 |
| TNF-α | Forward: 5’- acggcatggatctcaaagac -3’  Reverse: 5’- gtgggtgaggagcacgtagt -3’ |
| Jun | Forward: 5’-atgggcacatcaccactaca-3’  Reverse: 5’-gacactgggaagcgtgttct-3’ |
| Junb | Forward: 5’- gacgacctgcacaagatgaa-3’  Reverse: 5’- tgctgaggttggtgtagacg-3’ |
| Jund | Forward: 5’- gcctggaggagaaagtcaag-3’  Reverse: 5’- gttgacgtggctgaggactt-3’ |
| H2-Ab1 | Forward: 5’- acccagccaagatcaaagtg -3’  Reverse: 5’- atctccagcatgaccaggac -3’ |
| NFκB | Forward: 5’- cacctagctgccaaagaagg-3’  Reverse: 5’- gcaggctattgctcatcaca -3’ |
| GAPDH | Forward: 5’- tgatgggtgtgaaccacgag-3’  Reverse: 5’- gcccttccacaatgccaaag -3’ |
